# Profiling of Vaginal Microbial Communities in Chilean Women Via Self-sampling and Full-length 16S rRNA Sequencing

**DOI:** 10.1101/2025.03.21.644639

**Authors:** Barbara Oliva, Pablo Villanueva, Juan A. Ugalde

## Abstract

The vaginal microbiota is a dynamic ecosystem that plays a crucial role in women’s health, with Lactobacillus species predominating in healthy individuals. Disruptions in this microbiota can lead to dysbiosis, increasing susceptibility to infections such as bacterial vaginosis and other reproductive health complications. While most studies on the vaginal microbiome have been conducted in North American and European populations, data from Latin America, particularly Chile, remain scarce. This study presents a comprehensive analysis of the vaginal microbiota in Chilean women using full-length 16S rRNA gene sequencing with Oxford Nanopore Technology (ONT). We implemented a self-sampling methodology and a bioinformatics pipeline to achieve species-level resolution of microbial communities. Taxonomic profiling using Emu revealed that Lactobacillus was the dominant genus in most samples, while others exhibited microbial compositions consistent with Community State Type (CST) IV, characterized by higher diversity and lower Lactobacillus abundance. Alpha and beta diversity analyses further demonstrated distinct microbial clustering patterns across CSTs, with species-level resolution providing enhanced classification accuracy. Our findings highlight the potential of long-read sequencing and self-sampling for improving vaginal microbiome studies, particularly in underrepresented populations. This research contributes valuable insights into the composition and diversity of the vaginal microbiota in Chilean women, paving the way for more precise diagnostics and targeted interventions in reproductive health.

## Introduction

The vaginal microbiota is a dynamic and complex ecosystem composed of diverse microbial species that directly influence women’s overall and reproductive health. This microbial environment, predominantly dominated by bacteria of the *Lactobacillus* genus, plays a crucial role in protecting the epithelial mucosa of the vaginal tract by maintaining an acidic environment (pH 3.3-4.5), which inhibits the growth of pathogens and prevents viral infections ^1^. The balance between commensal and pathogenic species is essential to maintaining a healthy state, preventing infections and other gynecological complications.

*Lactobacillus* species metabolize glycogen from vaginal epithelial cells, promoting their proliferation and the production of lactic acid, which is vital for sustaining an acidic environment ^2^. Furthermore, these bacterial species secrete antimicrobial compounds, including hydrogen peroxide and bacteriocins, that further enhance vaginal defense mechanisms ^3^. The most frequently reported *Lactobacillus* species include *L. crispatus*, *L. gasseri*, *L. jensenii* and *L. iners* ^4^.

When the normal composition of the vaginal microbiota is disrupted, a dysbiotic state may develop, increasing vulnerability to infections such as bacterial vaginosis (BV) ^5^, vulvovaginal candidiasis ^6^, and sexually transmitted infections including chlamydia and HIV ^7,8^. Among these, BV is the most common, characterized by decreased *Lactobacillus* abundance and an increase in the abundance of anaerobic bacteria, such as *Gardenerella vaginalis*, *Atopobium vaginae*, and *Prevotella* species ^9^. Although BV can manifest with symptoms like malodorous vaginal discharge and itching, many women remain asymptomatic. However, this condition is associated with an increased risk of more severe infections and even complications during pregnancy ^10^.

Traditionally, clinical diagnosis of vaginal dysbiosis relies on conventional methods, including Amsel’s criteria, which assess clinical symptoms like elevated vaginal pH and microscopy evaluations ^11^, and the Nugent method, which uses Gram staining to bacterial species ^12^. However, these techniques are limited as they rely on human interpretation and do not provide an adequate resolution for detailed bacterial identification.

In recent years, advances in sequencing technologies have allowed us to overcome these limitations. 16S rRNA gene sequencing has become a powerful tool for studying complex microbial communities, as it enables bacterial identification without the need for cultures ^13^. However, most of the studies have been conducted using short-read sequencing platforms that do not span the entire 16S rRNA gene, therefore limiting species-level resolution ^14^. The introduction of long-read sequencing platforms, such as Oxford Nanopore (ONT), has opened new possibilities for studying the vaginal microbiota. ONT allows for long reads, facilitating the complete sequencing of the 16S rRNA gene (1,500 bp) and thus enabling species-level identification ^15^.

Beyond sequencing technology, accurate microbial classification also depends on the choice of reference databases. For example, the SILVA ^16^ database is widely used for taxonomic assignment in 16S rRNA gene studies, offering high-quality, curated sequences with detailed annotations. Meanwhile, the Genome Taxonomy Database (GTDB)^17^ provides a standardized classification based on genome phylogeny, making it particularly valuable for characterizing uncultivated or poorly defined microbial taxa. The selection of an appropriate database plays a crucial role in vaginal microbiota research, as it directly influences the accuracy of taxonomic assignments and, ultimately, our understanding of microbial communities and their impact on women’s health.

Microbiome studies using short-read sequencing approaches have classified the vaginal microbiota into distinct *Community State Types* (CST). CST I, II, III, and V are dominated by *Lactobacillus* species (*L. crispatus*, *L. gasseri*, *L. iners*, and *L. jensenii*, respectively), typically associated with healthy vaginal conditions. In contrast, CST IV lacks a clear dominant species and features a diverse range of anaerobic bacteria, such as *Prevotella*, *Dialister*, *Atopobium*, *Gardnerella*, *Megasphaera*, *Peptoniphilus*, *Sneathia*, *Eggerthella*, *Aerococcus*, *Finegoldia*, and *Mobiluncus*, often correlating with dysbiotic states ^10^.

Despite significant advancements in microbial profiling technologies, most vaginal microbiota studies have focused on North American and European populations, leaving substantial knowledge gaps for other populations, such as Latin America. In particular in Chile, studies on the vaginal microbiota are scarce and have primarily relied on standard clinical methods like the Amsel criteria. A study conducted with women attending a family health center, most of whom did not exhibit symptoms, concluded that 32% of the tested women had bacterial vaginosis (BV) ^18^. Similarly, a subsequent study using wet mount and Gram stain analysis found that 46.5% of women tested had some type of vaginal infection, with BV (16.8%) and vulvovaginal candidiasis (11.9%) being the most frequent infections ^19^. Both of these studies underscore the high prevalence of vaginal infections in Chilean women, but are limited by their reliance on conventional diagnostic methods. Given the limitations of these diagnostic approaches, there is an urgent need to study and understand the specific characteristics of vaginal microbial communities in Chilean women. Utilizing advanced microbial profiling techniques can provide a more comprehensive and detailed view of the microbial landscape. Understanding these microbial dynamics in a Chilean context is essential for improving diagnosis, prevention, and treatment strategies tailored to the specific needs of this population. Moreover, such research could reveal region-specific microbial profiles and health risks, which could have broader implications for public health and clinical management.

To address the need for more comprehensive profiling of vaginal microbiota in Chilean women, this study presents the development and application of a self-sampling method for vaginal swabs, along with a customized bioinformatic pipeline for estimating bacterial relative abundance. By performing full-length 16S rRNA gene sequencing using the ONT platform, we were able to obtain a species-level resolution taxonomic profile of the vaginal microbiome, identify species abundance, and classify samples into CSTs based on the microbiome composition. These results highlight the potential of self-sampling and long-read sequencing for expanding our understanding of vaginal health, particularly in underrepresented populations.

## Methods

### Implementation of a Self-Sampling Method for Vaginal Samples

#### Criteria and Study Cohort

This study was approved by the Bioethics Committee of the Faculty of Life Sciences, Universidad Andrés Bello. Twenty participants were recruited based on predefined criteria. Women between the ages of 22 and 36 who were able to provide informed consent were eligible for inclusion. Participants were excluded if they were menstruating at the time of sampling, pregnant, or had used antibiotics in the past 30 days. Additionally, all participants completed a detailed questionnaire covering demographic information, employment status, health habits, dietary patterns, hygiene practices, antibiotic use, and sexual history (Supplementary Table 1)

#### Vaginal Self-Sampling Protocol

The collection procedure was explained in detail to participants to ensure the consistency and reliability of the samples. Each participant was individually instructed to wash their hands before the procedure and perform an external vaginal cleansing using toilet paper moistened with water, wiping from the vulva towards the anus to prevent cross-contamination. For self-collection, participants were instructed to use the sterile swabs using the DNA/RNA Shield Collection Tube with Swab (Zymo Research, California, USA). Participants were asked to open the sterile container and hold the swab by the marked indentation. Using their free hand, they were instructed to gently separate the skin folds around the vaginal opening and insert the swab approximately 5 cm into the vaginal canal. They were then instructed to rotate the swab 10 times to ensure adequate sample collection and carefully withdraw it, avoiding contact with external surfaces. Immediately after collection, they were asked to place the swab into the tube containing the Shield buffer and store it at 4 °C until further processing.

### DNA Extraction and 16S Gene Sequencing

#### Genomic DNA Extraction and Library Preparation

DNA was extracted from the self-collected vaginal samples using the Quick-DNA Miniprep Plus Kit (Zymo Research, California, USA), following the manufacturer’s instructions. After a 2-minute incubation at room temperature, DNA was eluted in 50 µL of nuclease-free water. Concentration was measured using fluorometry (Qubit 4.0, Thermo Fisher Scientific, USA), while purity was determined by A260/A280 ratio using a Synergy spectrophotometer (Agilent, Biotek, California, USA). Libraries were prepared using the 16S Barcoding Kit (SQK-16S024, Oxford Nanopore Technologies, UK), designed to amplify the full-length 16S rRNA gene (1500 bp). Amplification was performed with genomic DNA (30 ng), universal primers 27F: 5’-AGAGAGTTTGATCMTGGCTCAG-3’ and 1492R: 5’-CGGTTACCTTGTTACGACTT-3’), which are flanked by additional sequences (barcodes) that differ for each sample and LongAmp Hot Start Taq Master Mix (New England Biolabs, USA). PCR conditions are detailed in Table 1.

**Table 1.**
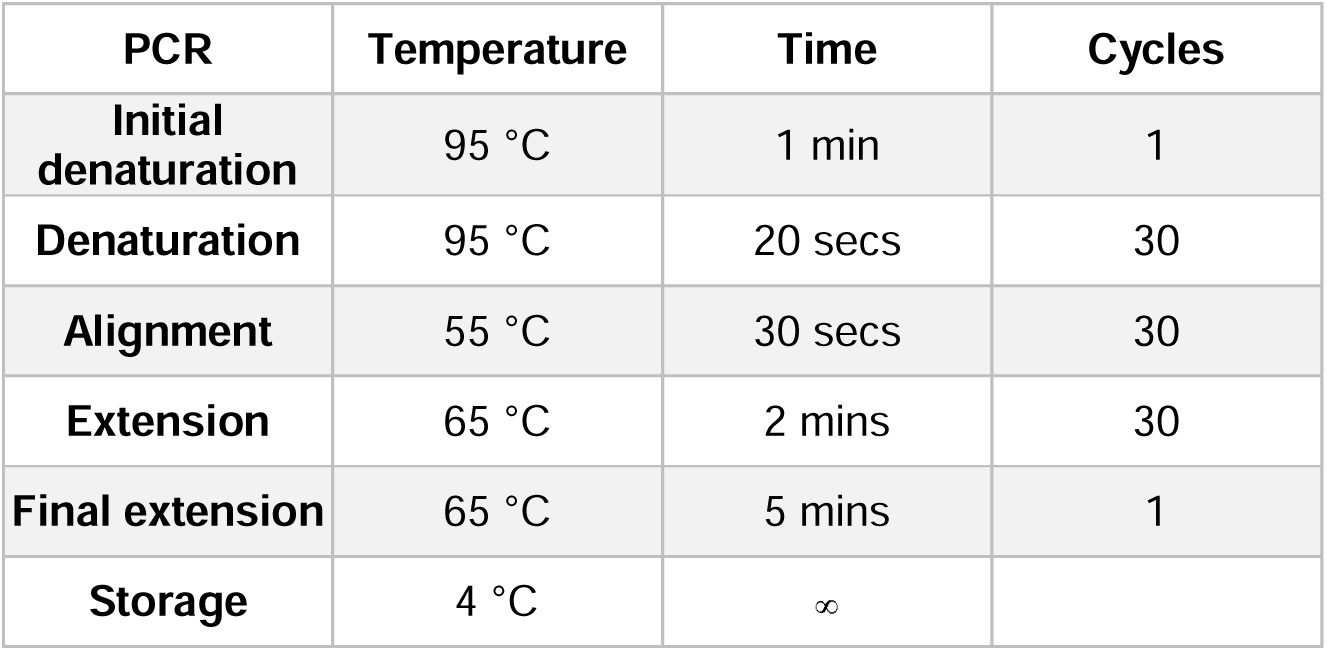
Conditions of the PCR cycles to amplify the bacterial 16S rRNA gene.

| PCR | Temperature | Time | Cycles |
| --- | --- | --- | --- |
| Initial denaturation | 95 °C | 1 min | 1 |
| Denaturation | 95 °C | 20 secs | 30 |
| Alignment | 55 °C | 30 secs | 30 |
| Extension | 65 °C | 2 mins | 30 |
| Final extension | 65 °C | 5 mins | 1 |
| Storage | 4 °C | ∞ |  |

Each sample was transferred to a 1.5 ml DNA LoBind Axygen tube and purified individually before pooling. Magnetic beads (Genosur, Chile) were resuspended by vortexing, and 30 μl were added to each sample and mixed by pipetting. After a 5 minute incubation on an orbital shaker, two washes were performed with 80% ethanol, carefully removing the supernatant without disturbing the pellet. The beads were then resuspended in 10 μl of 10 mM Tris-HCl, 50 mM NaCl, and incubated for 2 minutes at room temperature. A total of 10 μl of eluate was recovered, and the beads were discarded.

Libraries were quantified using fluorometry (Qubit 4.0, Thermo Fisher Scientific, USA). Barcoded libraries were pooled to a final concentration of 70 fmoles in 10 μl of 10 mM Tris-HCl, 50 mM NaCl (pH 8). Next, 1 μl of RAP was added, mixed gently, and incubated for 5 minutes at room temperature. The prepared and purified libraries were incubated on ice until loading into the flow cell.

#### 16S Gene Sequencing

The prepared libraries were mixed with 15 µL of the Sequencing Buffer (SB II), 10 µl Loading Beads (LB II), and nuclease-free water. The final adjusted sample was loaded into the flow cell R.9.4.1 (FLO-FLG001) and sequenced on the MinION Mk1B device (ONT, UK), which was connected to a desktop computer (Ryzen 9, 32 Gb RAM, RTX 3060). In this study, 10 samples were loaded at a time, and each sequencing session lasted about 24 hours. DNA sequencing data were acquired in POD5 format using the MINKNOW (version 24.06.16) software. The sequencing data for each sample were deposited in the NCBI SRA repository under BioProject code PRJNA1232418.

### Bioinformatic analysis

#### Sequenced Data Processing

Dorado (version 7.4.14) was used for basecalling and demultiplexing with the following settings: High accuracy model 450 bps, Barcode Kit SQK-PBK004, and trim_barcodes=on. The resulting sequence data were generated as FASTQ files. These files underwent length filtering using fastp ^20^, with trimming and quality filtering options disabled. Reads were retained if their lengths ranged between 1400 and 1700 base pairs (--length_required 1400 - -length_limit 1700) for expected full 16S rRNA gene length.

#### Vaginal mock community generation and analysis

A vaginal mock community was simulated for database comparisons using the genome sequences of some of the most common taxa reported in vaginal communities. Genome sequences were downloaded from the NCBI genome database ^21^, and details of each assembly are provided in Table 2. All genomes were concatenated and used as a reference genome for Grinder (version 0.5.3) ^22^ to simulate 16S amplicons. Standard full-length 16S primer sequences were applied for the forward_reverse parameter (27F: 5’-AGAGTTTGATYMTGGCTCAG-3’; 1492R: 5’-TACGGYTACCTTGTTACGACTT-3’), with 1000 reads specified for the total_reads parameter. The resulting error-prone reads were then processed using CuRe-Sim-LoRM ^23^ to introduce ONT-associated errors, using the tool default parameters.

**Table 2.** Genome sequences used for vaginal 16S mock community.

| Organism Scientific Name | Assembly Accession | Size |
| --- | --- | --- |
| *Ureaplasma parvum* ATCC 27815 | GCA_000019345.1 | 751,679 |
| *Prevotella bivia* DSM 20514 | GCA_000262545.1 | 2,521,238 |
| *Lactobacillus gasseri* 2016 | GCA_000439915.2 | 1,892,784 |
| *Streptococcus agalactiae* | GCA_001552035.1 | 2,079,123 |
| *Lactobacillus jensenii* | GCA_001936235.1 | 1,672,949 |
| *Gardnerella vaginalis* | GCA_002861965.1 | 1,675,681 |
| *Anaerococcus hydrogenalis* | GCA_002871895.1 | 1,908,330 |
| *Lactobacillus iners* | GCA_009556455.1 | 1,363,099 |
| *Lactobacillus crispatus* | GCA_009769205.1 | 2,039,404 |
| *Bifidobacterium breve* | GCA_024665435.1 | 2,336,700 |
| *Megasphaera paucivorans* | GCA_900103535.1 | 2,912,276 |

The obtained reads were analyzed using Emu (version 3.4.5) ^24^ with three databases. These included the default Emu database (version 3.4.5), which integrates rrnDB version 5.6 ^25^ and NCBI 16S RefSeq ^26^; the pre-built Silva database (version 138.1) ^16^; and a custom GTDB database adapted for Emu. The custom GTDB database was constructed using GTDB release 220 ^17^ and the Emu build-database option. Taxonomic assignments were performed using the Emu abundance function for each database. The relative abundance output tables were subsequently analyzed and compared against expected relative abundance values using R (version 4.2.2).

#### Taxonomic Profiling and Data Analysis

Filtered reads were analyzed per sample to determine the taxonomic relative abundance of 16S genomic sequences using Emu (version 3.4.5) ^14^. The analysis was conducted using the Emu abundance function, with all parameters set to default. Between the aforementioned databases, the default Emu database (version 3.4.5, rrnDB ^25^ and NCBI 16S RefSeq ^26^) was chosen based on its performance. The relative abundance output tables were imported into R and converted into phyloseq objects using the *phyloseq* package (version 1.42.0) ^27^. Taxonomic data were aggregated at the genus or species level using the tax_glom() function in phyloseq. Filtered and processed data were visualized using the *ggplot2* package (version 3.5.1) ^28^. Alpha diversity and beta diversity (Bray-Curtis PCoA distance) for each sample were calculated using the plot_richness() and ordinate() functions in the *phyloseq* package.

## Results

### *In Silico* Validation of 16S Metataxonomic Analysis

#### Vaginal Mock community and database selection

To select the most suitable database for this dataset, a comparison of different 16S taxonomic databases was conducted using an *in silico*-generated vaginal mock community. Following information from the literature ^10^, the mock community was designed to represent the most common bacterial taxa found in vaginal samples across all CSTs. A total of 1,000 *in silico* 16S amplicons were simulated using Grinder (version 0.5.3) ^22^ with its *in silico* PCR option, employing genomic DNA sequences from the selected taxa and standard full-length 16S primer sequences. From the 11 taxa initially selected, seven amplified successfully under these simulation conditions. The estimated relative abundances of the amplicons for each taxon are summarized in Table 3.

**Table 3.** Species present in in-silico mock community. All species are in a equal relative abundance of 12.5% in the mock community.

| Taxa | Amplified seq_id |
| --- | --- |
| *Prevotella bivia* | NZ_JH660658.1 |
| *Prevotella bivia* | NZ_JH660660.1 |
| *Lactobacillus crispatus* | NZ_CP039266.1 |
| *Lactobacillus jensenii* | NZ_CP018809.1 |
| ***Streptococcus agalactiae*** | NZ_CP012480.1 |
| ***Anaerococcus hydrogenalis*** | NZ_PNHP01000002.1 |
| ***Lactobacillus iners*** | NZ_CP045664.1 |
| ***Lactobacillus gasseri*** | NZ_CP090409.1 |

**Table 4.** Relative genus abundances of *in silico* vaginal mock community estimated with different databases.

| Genus | Expected | Silva | GTDB | EMU |
| --- | --- | --- | --- | --- |
| ***Anaerococcus*** | **0.125** | 0.140 | 0.141 | 0.140 |
| ***Lactobacillus*** | **0.5** | 0.488 | 0.488 | 0.494 |
| ***Prevotella*** | **0.25** | 0.233 | 0.232 | 0.232 |
| ***Streptococcus*** | **0.125** | 0.134 | 0.134 | 0.131 |
| ***Geobacillus*** | - | 0.001 | 0 | 0 |
| ***Limosilactobacillus*** | - | 0.002 | 0 | 0 |
| **UCG-004** | - | 0.001 | 0 | 0 |
| ***Akkermansia*** | - | 0 | 0.001 | 0 |
| ***Bifidobacterium*** | - | 0 | 0.002 | 0 |
| ***Ligilactobacillus*** | - | 0 | 0.001 | 0 |
| ***Bacillus*** | - | 0 | 0 | 0.001 |
| ***Lactiplantibacillus*** | - | 0 | 0 | 0.001 |
| **Total miss assigned** | - | 0.004 | 0.004 | 0.002 |

Error was introduced into the *in silico* simulated amplicons to mimic ONT long-read sequencing using CuReSim-LoRM ^23^. The resulting reads constituted the final vaginal mock community. Considering the benchmarking performance of the Emu tool ^14^, this method was selected for 16S taxonomic assignment. Three 16S databases were evaluated with this tool using the generated mock community: the Emu database, the SILVA database version available for Emu, and a GTDB database version adapted for Emu. Relative abundances obtained for the mock community at the genus level for each database are shown in Table 3 and graphically represented in Figure 1. The estimated relative abundances from all databases were close to the expected values, with minor deviations attributable to simulated sequencing errors or taxonomic misassignments. Among the three databases, the Emu database demonstrated the highest accuracy, showing relative abundances closest to the expected values and misassigning only two genera.

**Figure 1.**
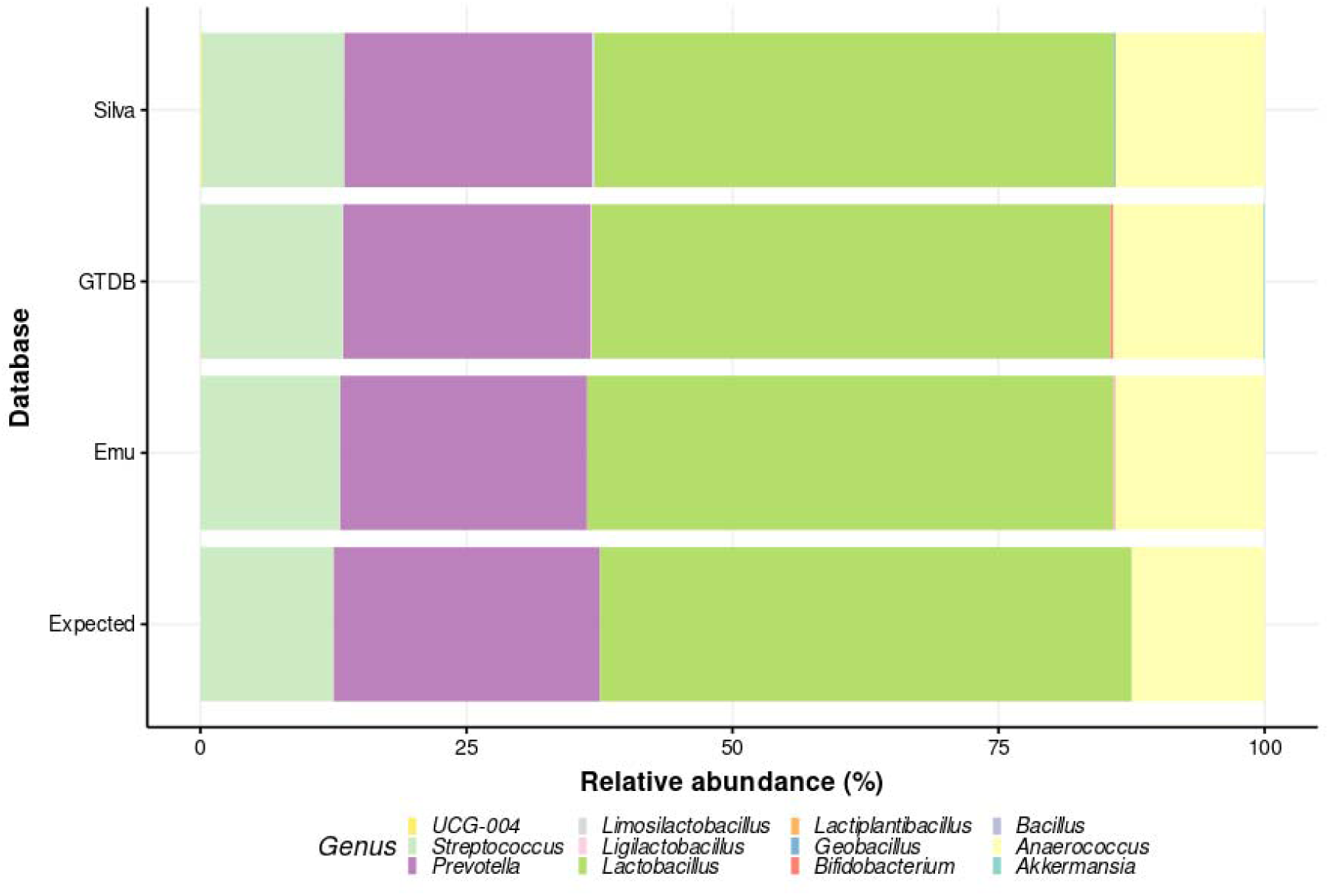
Relative abundance plot of the *in silico* vaginal mock community comparing the three databases used. Bars represent relative abundances estimated in the mock community with different databases using the Emu tool.

The same analysis was conducted at the species level for the three databases, with a summarized version of the results presented in Table 5. In all cases, the relative abundances closely align with the expected values; however, certain species are either underrepresented or overrepresented. For instance, *Lactobacillus crispatus* is consistently overrepresented across all databases, while *Prevotella bivia* is underrepresented. The SILVA database showed a significant limitation, with five taxa being assigned only at the genus level rather than resolving to the species level, a drawback not observed with the other databases. In contrast, the GTDB and Emu databases demonstrated better species-level resolution but exhibited a higher degree of misassigned species. Specifically, the GTDB database misassigned 14 species, while the Emu database misassigned 11 species, with the latter showing lower relative abundances for these misassignments. Considering the importance of resolving taxa at the species level for subsequent analyses and minimizing taxonomic assignment errors in data comparable to the generated mock community, the Emu database was selected for further analyses.

**Table 5.**
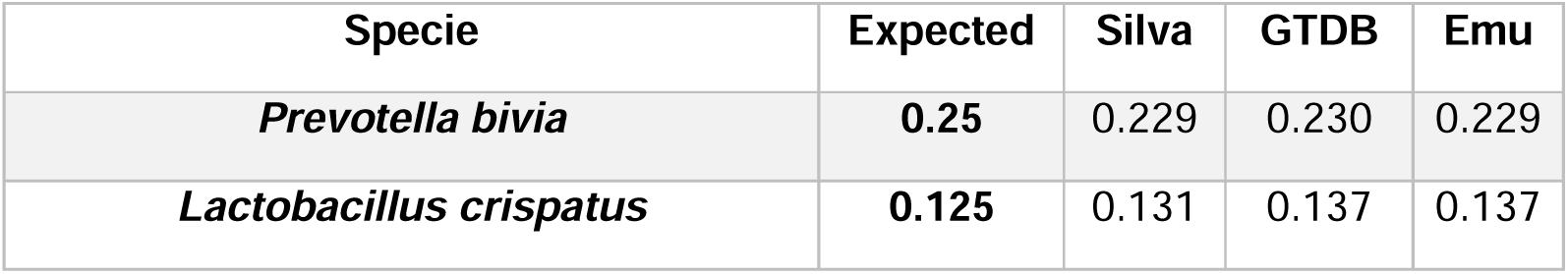
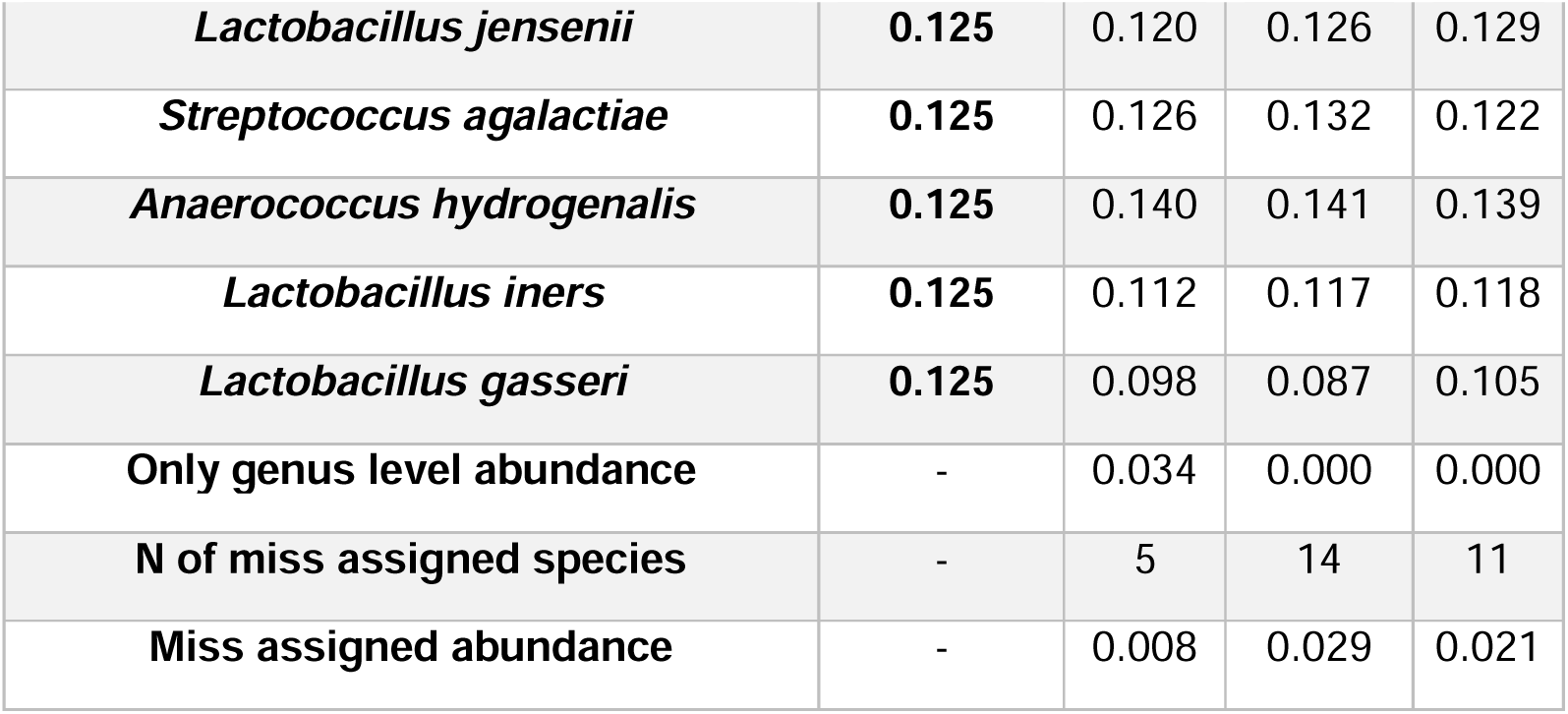
Relative species abundances of *in silico* vaginal mock community estimated with different databases.

### ONT Sequencing and 16S Metataxonomic Analysis of Self-Collected Vaginal Samples

#### Characteristics of participants

In this study, 20 women residing in Chile between the ages of 22 and 36 years (median age 32 years) consented to vaginal self-sampling. Each participant responded to an interview related to her lifestyle and health history. All participants declared themselves sexually active. Among them, 6 out of 20 reported having recurrent infections at least three times a year. In addition, 25% of the participants reported using antibiotics regularly. Regarding hygienic practices, responses varied, but mainly water use and 6 participants reported daily use of intimate hygiene products. During the self-sampling procedure, only one participant experienced discomfort when inserting the swab.

#### MinION-Based 16S rRNA Sequencing and Taxonomic Profiling of Vaginal Microbiota

To characterize the bacterial composition of vaginal samples, DNA extraction was performed, yielding concentrations ranging from 30 to 223 ng/µL, except for one sample with an insufficient concentration (<0.2 ng/µL). Library preparation and purification were carried out for all extracted samples, which were successfully sequenced using the Oxford Nanopore Technologies (ONT) MinION platform. Sequencing and basecalling results yielded an average read length of 1.6 Kb. For taxonomic profiling, the 20 barcoded samples were length-filtered using fastp ^20^, retaining only reads between 1,400 and 1,700 base pairs, corresponding to the estimated full-length 16S rRNA gene. The number of reads obtained per sample and read counts after filtering are summarized in Table 6. On average, 7.7% (SD = 1.9%) of reads per sample were retained after filtering.

**Table 6.** Reads for each vaginal sample after sequencing and length filtering.

| **Sample** | **Reads Before Filtering** | **Reads After Filtering** | **filtered%** |
| --- | --- | --- | --- |
| **M1** | 23,343 | 21,698 | 7.05 |
| **M2** | 20,455 | 19,116 | 6.55 |
| **M3** | 27,351 | 25,735 | 5.91 |
| **M4** | 34,603 | 32,543 | 5.95 |
| **M5** | 40,313 | 37,919 | 5.94 |
| **M6** | 11,000 | 10,246 | 6.85 |
| **M7** | 8,000 | 7,444 | 6.95 |
| **M8** | 5,000 | 4,596 | 8.08 |
| **M9** | 9,000 | 8,359 | 7.12 |
| **M10** | 5,000 | 4,663 | 6.74 |
| **M11** | 38,000 | 35,394 | 6.86 |
| **M12** | 9,000 | 8,360 | 7.11 |
| **M13** | 6,000 | 5,542 | 7.63 |
| **M14** | 10,000 | 9,372 | 6.28 |
| **M15** | 14,000 | 13,003 | 7.12 |
| **M16** | 13,000 | 12,142 | 6.60 |
| **M17** | 12,263 | 11,060 | 9.81 |
| **M18** | 15,757 | 13,760 | 12.67 |
| **M19** | 17,096 | 15,261 | 10.73 |
| **M20** | 42,993 | 37,830 | 12.01 |

Filtered reads from each sample were taxonomically profiled using Emu ^14^ and the Emu 16S database. Notably, there were no unassigned reads across all samples, indicating that the Emu pipeline successfully classified all 16S genes present. The relative abundances obtained were processed in R as a phyloseq object for downstream analysis and visualization. The estimated abundances per sample at the genus level are presented in Figure 2, with the complete dataset available in Supplementary Table 2. As expected, the genus *Lactobacillus* was the most abundant across samples, with 14 out of 20 samples showing a relative abundance above 75%, 12 exceeding 90%, and 5 surpassing 99%.

**Figure 2.**
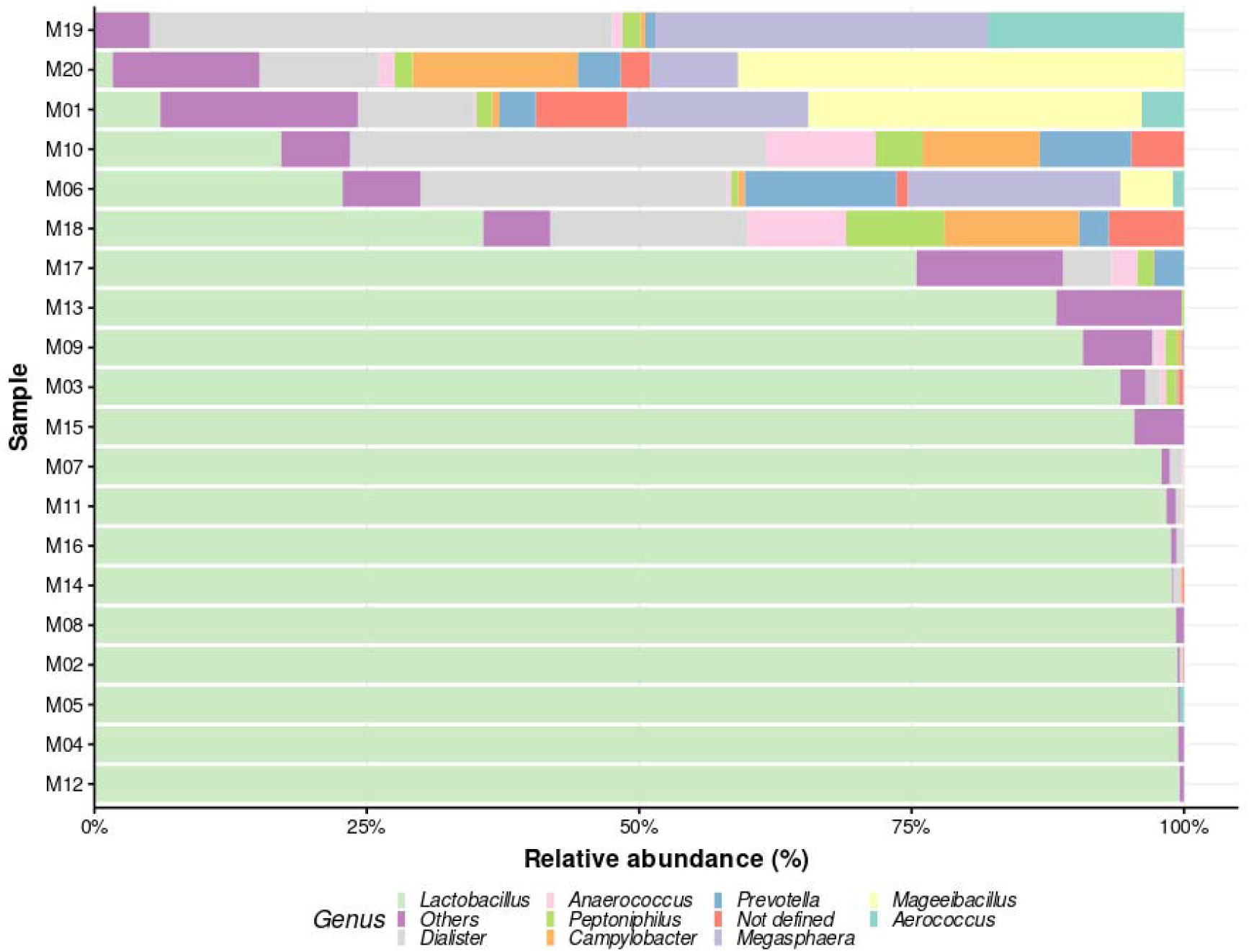
Bar plot of the top ten most abundant genera across all analyzed vaginal 16S samples. The bars represent relative abundances estimated by Emu for each sequenced sample, displaying only the top ten genera. Remaining genera are grouped as “Others.” The x-axis indicates the percentage of estimated abundances, while the y-axis lists samples ordered ascendingly by *Lactobacillus* abundance. The “Not defined” category corresponds to a particular taxon (*Peptostreptococcaceae* bacterium oral taxon 929) that lacks an assigned genus in the database used for analysis.

The second most abundant classification was the “Others” category, encompassing less represented genera that did not rank among the top 10. This pattern suggests that in samples dominated by *Lactobacillus*, the less abundant genera are a mix of low-abundance taxa without a consistent secondary dominant genus. Conversely, in the six samples where *Lactobacillus* relative abundance fell below 50%, the second most abundant, and in some cases, the dominant, genus was *Dialister*. This genus, along with others evenly represented in these samples, aligns with the composition of Community State Type (CST) IV, characterized by the absence of a dominant genus ^29^.

Relative abundances were analyzed at the species level, with results presented in Figure 3, while the complete dataset is available in Supplementary Table 3. Among the *Lactobacillus* species, *L. crispatus*, *L. iners*, and *L. gaseri* were the most relatively abundant, correlating strongly with community state types (CST) I, II, and III, respectively ^10^. These species exhibited relative abundances often exceeding 90% in dominant samples, such as M07, M12, and M14. Notably, 7 out of the 20 samples exhibited a relative abundance of *L. crispatus* above 80%, aligning with healthy vaginal microbiota profiles. Interestingly, only one sample (M11) showed a dominant abundance of *L. jensenii* (98%), which is characteristic of CST V. It is also noteworthy that in cases where *L. iners* was the most abundant species (M05, M09, M13), its relative abundance never exceeded 80%, and the second most abundant species was also a *Lactobacillus* species.

**Figure 3.**
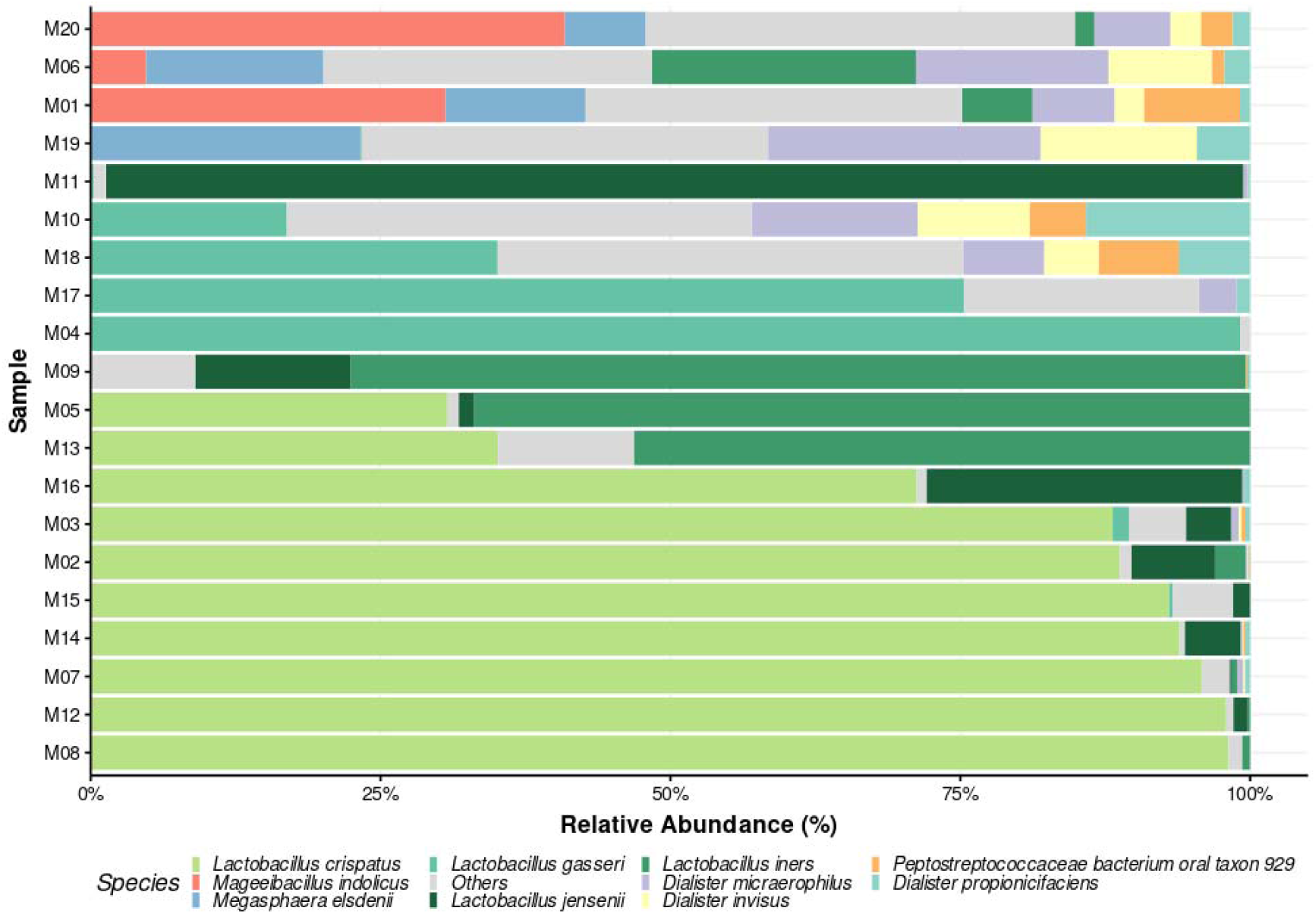
Bar plot of the top ten most abundant species across all analyzed vaginal 16S samples. The bars represent the species relative abundances estimated by Emu for each sequenced sample, displaying only the top ten most abundant across all samples. Remaining species are grouped as “Others.” The x-axis indicates the percentage of estimated abundances, while the y-axis lists samples ordered ascendingly by species from the *Lactobacillus* genus abundance.

Samples with reduced *Lactobacillus* dominance, such as M01 and M19, displayed more diverse microbial communities, including significant proportions of *Mageeibacillus indolicus* (30.6% in M01) and *Megasphaera elsdenii* (23.2% in M19). The “Others” category, representing less abundant taxa, contributed up to 30-40% in the more diverse samples (M01, M06, M10, M18, M19, M20), indicating substantial microbial complexity. In particular, samples M19 and M20, where *Lactobacillus* species were minimal, exhibited a marked increase in diversity, with dominance by genera such as *Mageeibacillus* and *Megasphaera*. This shift toward increased diversity and reduced *Lactobacillus* dominance is consistent with CST IV profiles.

Based on these results, the samples were preliminarily classified into different CSTs based on the most relatively abundant *Lactobacillus* species. Dominance was defined as a relative abundance of a species exceeding 80%. The results are presented in table 7.

**Table 7.** Classification of Samples into Community State Types (CSTs) Based on Relative Abundances of *Lactobacillus* Species.

| **Sample** | **Most abundant Specie** | **Abundance (%)** | **CST** | **Notes** |
| --- | --- | --- | --- | --- |
| **M01** | *Mageeibacillus indolicus* | 30.60% | IV | Low *Lactobacillus*, diverse anaerobes |
| **M02** | *L. crispatus* | 88.70% | I | *L. crispatus* dominant |
| **M03** | *L. crispatus* | 88.10% | I | *L. crispatus* dominant |
| **M04** | *L. gasseri* | 99.20% | II | *L. gasseri* dominant |
| **M05** | *L. iners* | 66.90% | III | *L. iners* mainly dominant (*L. crispatus* second most abundant) |
| **M06** | Diverse (others) | 28.30% | IV | Low *Lactobacillus*, diverse anaerobes |
| **M07** | *L. crispatus* | 95.80% | I | *L. crispatus* dominant |
| **M08** | *L. crispatus* | 98.10% | I | *L. crispatus* dominant |
| **M09** | *L. iners* | 77.20% | III | *L. iners* mainly dominant (*L. jensenii* second most abundant) |
| **M10** | Diverse (others) | 40% | IV | Low *Lactobacillus*, diverse anaerobes |
| **M11** | *L. jensenii* | 98% | V | *L. jensenii* dominant |
| **M12** | *L. crispatus* | 97.90% | I | *L. crispatus* dominant |
| **M13** | *L. iners* | 53.10% | III | *L. iners* mainly dominant (*L. crispatus* second most abundant) |
| **M14** | *L. crispatus* | 93.90% | I | *L. crispatus* dominant |
| **M15** | *L. crispatus* | 93.00% | I | *L. crispatus* dominant |
| **M16** | *L. crispatus* | 71.20% | I | *L. crispatus* mainly dominant (*L. jensenii* second most abundant) |
| **M17** | *L. gasseri* | 75.30% | II | *L. gasseri* mainly dominant (Others second most abundant) |
| **M18** | Diverse (others) | 40% | IV | Low *Lactobacillus*, diverse anaerobes |
| **M19** | *Mageeibacillus indolicus* | 35.00% | IV | Low *Lactobacillus*, diverse anaerobes |
| **M20** | *Mageeibacillus indolicus* | 40.90% | IV | Low *Lactobacillus*, diverse anaerobes |

### Diversity analysis of vaginal microbiome samples

#### Genus diversity analysis correlates with potential dysbiosis state

After classifying each sample into a community state type (CST), we conducted a diversity analysis at the genus level to evaluate the information provided at this taxonomic resolution and compare it with species-level resolution. This comparison aims to highlight the advantages of full-length 16S gene sequencing. Alpha diversity, measured using the Shannon and Simpson indices, was calculated for each sample at the genus level. The results are presented in Figure 4, with statistical comparisons of the diversity indices across CSTs shown in Table 8. As expected, samples classified as CST IV exhibited the highest diversity among all groups. Although the sample size is limited, CST I had the lowest mean diversity, suggesting that this state is predominantly dominated by *Lactobacillus* with minimal presence of other bacterial genera. Given that CST IV is often associated with a potential dysbiotic state, its significantly higher mean diversity suggests that alpha diversity at the genus level may serve as an indicator of dysbiosis when considered independently

**Figure 4.**
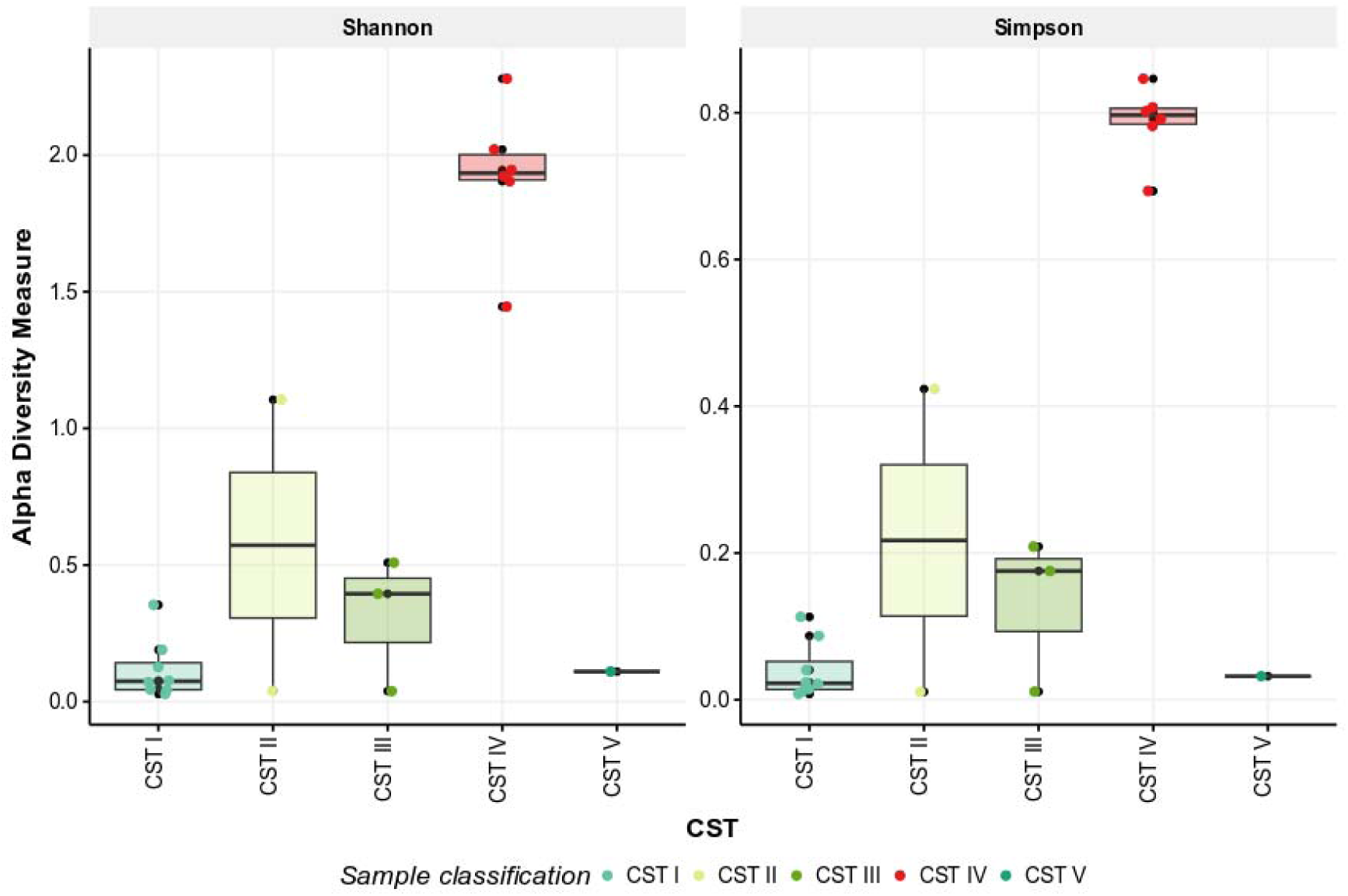
Alpha diversity indices (Shannon and Simpson) calculated at the bacterial genus level in vaginal samples. Boxplots represent genus-level diversity in vaginal samples, grouped according to previously determined community state types (CSTs). CSTs include: CST I (*L. crispatus*-dominant), CST II (*L. gasseri*-dominant), CST III (*L. iners*-dominant), CST IV (high bacterial diversity), and CST V (*L. jensenii*-dominant). The x-axis displays the different CSTs, where each point represents an individual sample. The y-axis indicates alpha diversity, measured using Shannon and Simpson indices.

**Table 8.** Alpha diversity statistics analysis at genus level grouped by community states.

| **CST** | **Mean Shannon** | **SD Shannon** | **Mean Simpson** | **SD Simpson** |
| --- | --- | --- | --- | --- |
| **CST I** | 0.117 | 0.109 | 0.040 | 0.039 |
| **CST II** | 0.573 | 0.753 | 0.217 | 0.292 |
| **CST III** | 0.314 | 0.245 | 0.132 | 0.106 |
| **CST IV** | 1.920 | 0.270 | 0.787 | 0.051 |
| **CST V** | 0.110 | - | 0.032 | - |

A similar analysis was performed for beta diversity at the genus level using Bray-Curtis distance, with results shown in Figure 5. Principal component analysis results are provided in Supplementary Figure 4. Similar to the alpha diversity analysis, all samples grouped according to their previously assigned community state type, except for those classified as CST IV. These results indicate that, at the genus level, it is not possible to distinguish between the *Lactobacillus*-dominated community states, but samples corresponding to CST IV can be clearly separated. In Figure 5, the principal component on the X-axis explains 75.2% of the variance, demonstrating that this type of analysis at the genus level provides a strong statistical representation. The loading analysis further indicates that the primary contributors to this separation are the relative abundances of *Lactobacillus* and various other bacterial genera.

**Figure 5.**
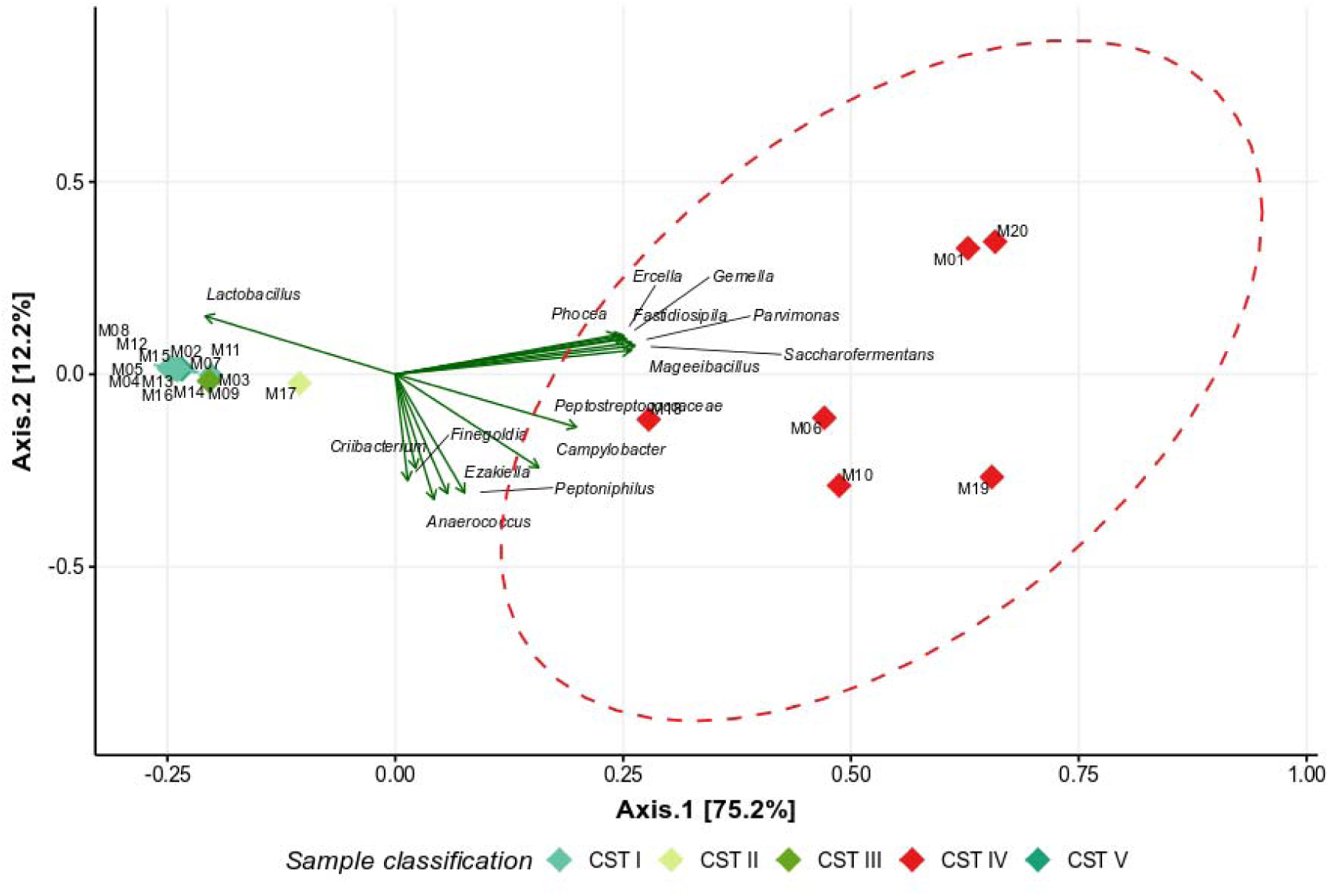
Beta Diversity Analysis by PCoA Based on Bray-Curtis Distance at the Genus Level. The graph represents the Bray-Curtis distances of vaginal samples analyzed at the genus level. Each sample is color-coded according to its previously determined community state. All samples classified as CST IV were grouped to add an ellipse (red dashed line). PCA loadings are represented as arrows labeled with the corresponding genera, displaying only the 15 most contributing taxa. The X-axis represents the first principal component, while the Y-axis represents the second principal component.

#### Species level diversity analysis allows community state classification

To further assess taxonomic resolution in our samples, we conducted a diversity analysis at the species level. Alpha diversity, grouped by CSTs, is shown in Figure 6, with statistical comparisons presented in Table 9. As observed at the genus level, diversity indices varied significantly across vaginal community states. CST IV exhibited the highest diversity values, with an average Shannon index of 2.42 and a Simpson index of 0.851, indicating a markedly more heterogeneous bacterial composition. Conversely, CSTs dominated by *Lactobacillus* species (CST I, II, III, and V) displayed lower diversity values, reinforcing the notion of a more uniform microbiota. These results further support the use of alpha diversity as a numerical indicator of potentially dysbiotic states, as CST IV consistently showed elevated diversity compared to *Lactobacillus*-dominated CSTs. However, unlike the genus-level analysis, the species-level resolution suggests potential for CST classification, as mean diversity values appear to distinguish each CST more clearly. Although limited by sample size, these findings highlight the added value of species-level analysis in refining community state characterization.

**Figure 6.**
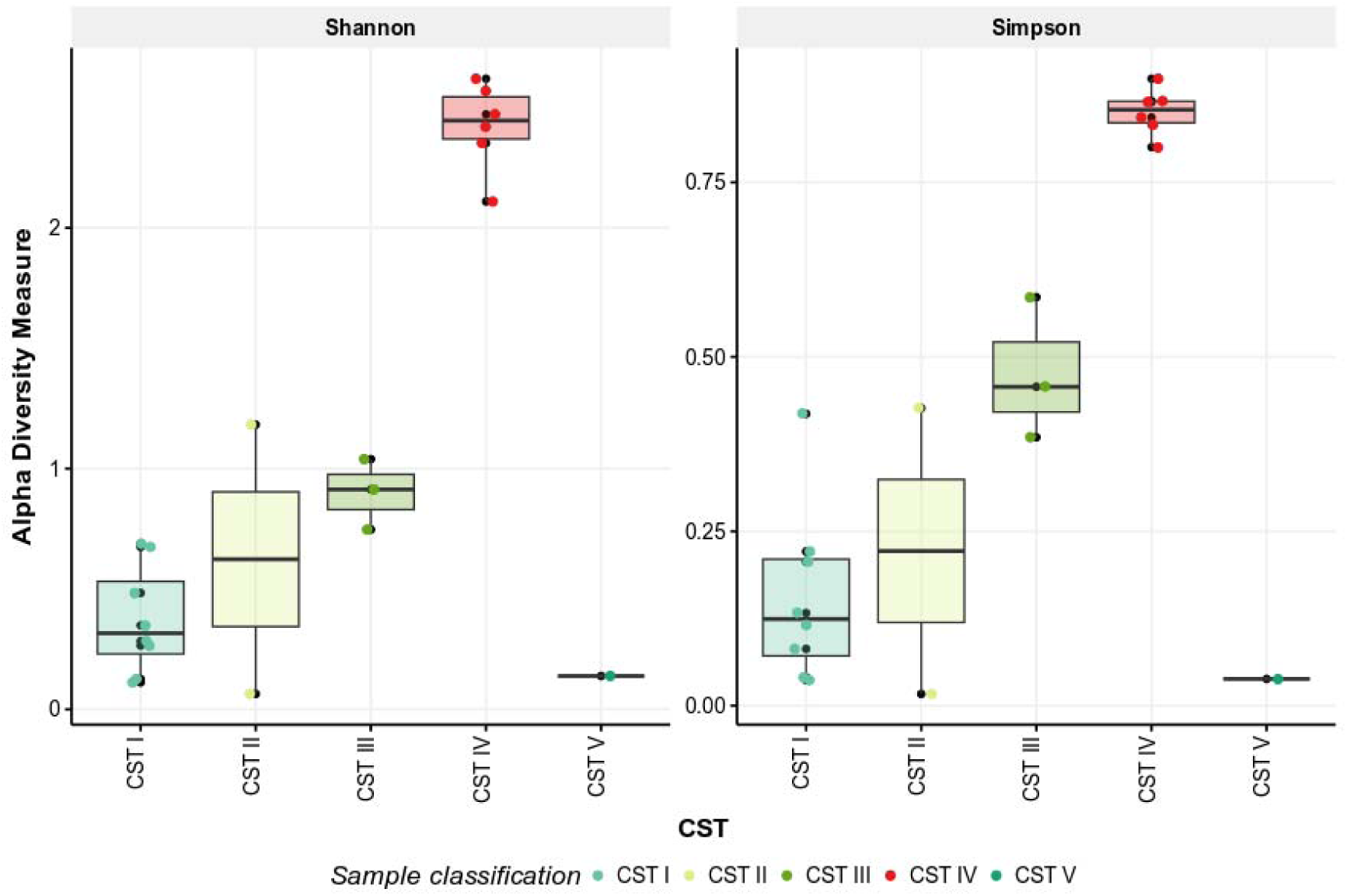
Alpha Diversity Analysis at the Species Level in Vaginal Samples. Boxplots represent species-level diversity in vaginal samples, grouped according to community state types (CSTs). CSTs include: CST I (*L. crispatus*-dominant), CST II (*L. gasseri*-dominant), CST III (*L. iners*-dominant), CST IV (high bacterial diversity), and CST V (*L. jensenii*-dominant). The x-axis displays the different CSTs, where each point represents an individual sample. The y-axis indicates alpha diversity, measured using Shannon and Simpson indices.

**Table 9.** Alpha diversity statistics analysis at species level grouped by community states.

| **CST** | **Mean_Shannon** | **SD_Shannon** | **Mean_Simpson** | **SD_Simpson** |
| --- | --- | --- | --- | --- |
| **CST I** | 0.373 | 0.224 | 0.157 | 0.126 |
| **CST II** | 0.624 | 0.791 | 0.222 | 0.290 |
| **CST III** | 0.900 | 0.147 | 0.476 | 0.102 |
| **CST IV** | 2.42 | 0.182 | 0.851 | 0.0336 |
| **CST V** | 0.138 | - | 0.0383 | - |

The same analysis was performed for beta diversity at the species level using Bray-Curtis distance, with results shown in Figure 7. Additional results are provided in Supplementary Table 6. Similar to the alpha diversity analysis, samples clustered according to their previously assigned community state type (CST), further validating the classification. However, unlike the genus-level analysis, beta diversity at the species level allowed for a clearer differentiation between all CSTs, indicating a finer resolution of microbial composition at this taxonomic level. In Figure 7, the first principal component (X-axis) explains 44% of the variance, demonstrating a more complex distribution of diversity compared to the genus-level analysis. Notably, CST IV exhibits the widest dispersion, reinforcing the idea that this community state encompasses a highly diverse and heterogeneous microbial composition. The observed variability suggests the possibility of sub-classifications within CST IV, potentially corresponding to different dysbiotic states or distinct ecological shifts within the vaginal microbiota.The loading analysis, highlighting the 15 most influential taxa, reveals that multiple species contribute significantly to the differentiation among CSTs. In particular, species associated with CST IV, such as *Neisseria gonorrhoeae*, *Prevotella corporis*, and *Campylobacter hominis*, contrast sharply with those predominant in *Lactobacillus*-dominated CSTs. This suggests that species-level resolution provides a more detailed understanding of the microbial landscape, capturing ecological variations that may be overlooked at the genus level. Despite the limited sample size, these results highlight the potential of species-level beta diversity analysis as a more precise tool for identifying microbial shifts, particularly in dysbiotic states. Future studies with larger datasets and longitudinal samples could further refine CST classification and explore its clinical implications.

**Figure 7.**
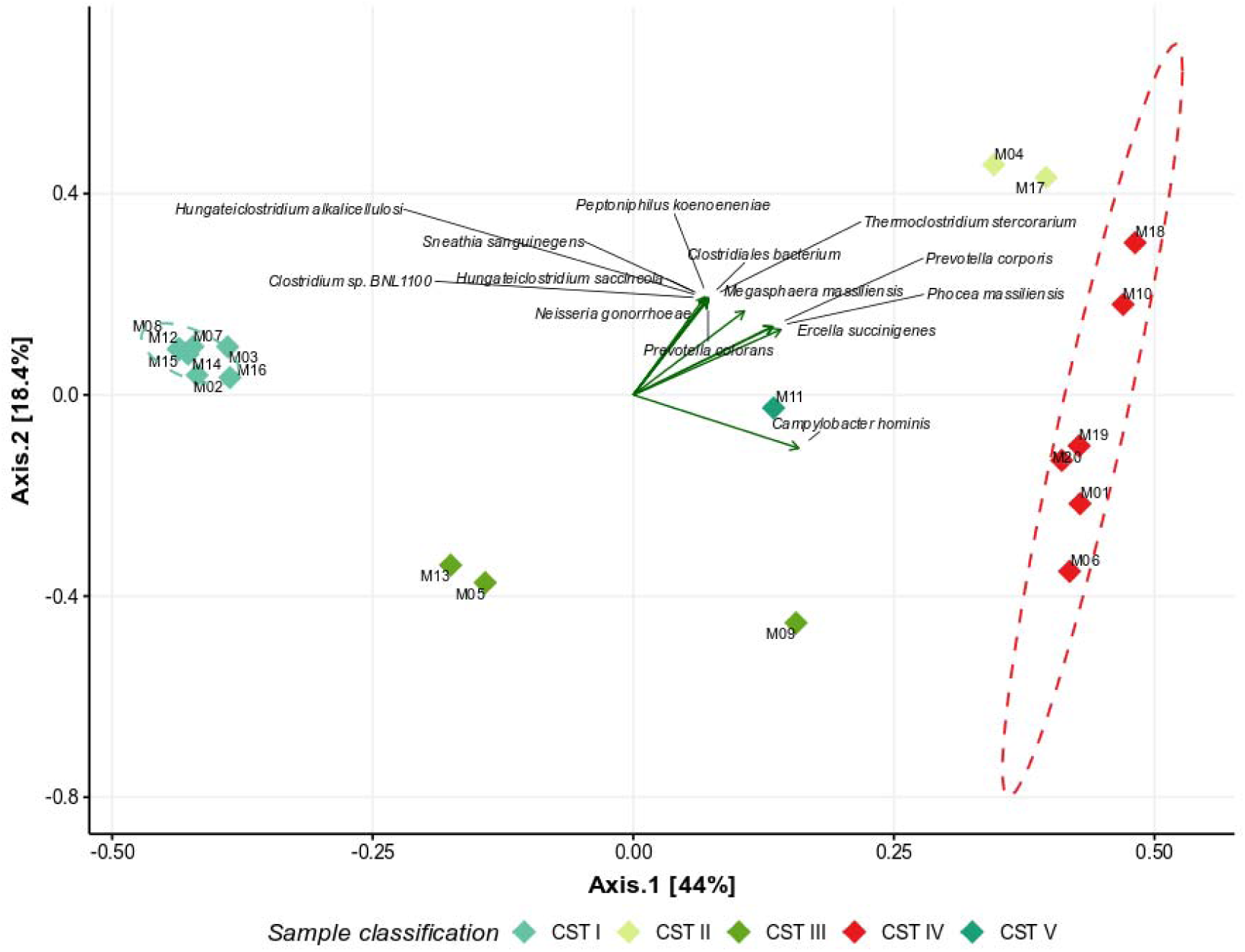
Beta Diversity Analysis by PCoA Based on Bray-Curtis Distance at the Species Level. The graph represents the Bray-Curtis distances of vaginal samples analyzed at the species level. Each sample is color-coded according to its previously determined community state (CST). Ellipses were added to group samples by CST; however, due to limited sample size, only CST I and CST VI are displayed. Principal component analysis (PCA) loadings are represented as arrows labeled with the corresponding species, showing only the 15 most contributing taxa. The x-axis represents the first principal component, while the y-axis represents the second principal component.

## Discussion

This study successfully characterized the vaginal microbial communities of 20 female residents in Chile using 16S rRNA gene sequencing with Oxford Nanopore Technologies (ONT). Our findings provide valuable insights into the taxonomic composition and Community State Types (CSTs) of vaginal microbiota ^10^, highlighting the applicability of long-read sequencing for vaginal microbial profiling. Importantly, this is the first study of its kind conducted in Chile, filling a critical gap in the understanding of vaginal microbiota within the region and establishing a baseline for future research in Latin America.

The implementation of ONT-based 16S sequencing demonstrated its potential for accurately characterizing the vaginal microbiota at both the genus and species levels, as highlighted by previous studies using MinION technology for rapid microbiota analysis in clinical settings ^30^. The *in silico* mock community validation confirmed the accuracy of different databases for taxonomic classification. Among the databases evaluated, Emu exhibited the highest accuracy ^24^, with minimal taxonomic misassignments compared to SILVA and GTDB. This reinforces the importance of selecting an appropriate reference database for taxonomic profiling, particularly in microbial communities where accurate species-level classification is critical for ecological and clinical interpretations.

The processing of vaginal samples, specifically in DNA extraction, variable concentrations were obtained, with one sample presenting an insufficient amount for downstream analysis, which underscores the importance of careful sample handling and optimization in low-biomass environments. The sequencing results demonstrated an average read length of 1.6 kb, ensuring near-full-length 16S rRNA coverage, which enhanced taxonomic resolution and allowed for a more accurate characterization of the microbial communities. This result is consistent with previous studies demonstrating that nanopore sequencing of complete 16S rRNA gene amplicons from human intestinal and vaginal microbiota provides further information to determine the bacterial species. ^30–32^

Our findings are consistent with established CST classifications, demonstrating that *Lactobacillus*-dominated communities (CST I, II, III and V) were prevalent in the sampled cohort. In particular, we observed that 14 of 20 participants had an abundance of *Lactobacillus* greater than 75%, with *L. crispatus* as the most predominant species. This finding supports previous studies that highlight the protective role of *L. crispatus* in vaginal health by promoting lactic acid production and maintaining a low vaginal pH, which contributes to preventing urogenital infections and bacterial vaginosis (BV) ^1^. Moreover, cohort studies from the United States, Europe and Asia have consistently reported a high prevalence of *L. crispatus* in women with a healthy vaginal microbiota, reinforcing its role in vaginal homeostasis and infection resistance ^10,33,34^.

Conversely, *Lactobacillus iners* was found in moderate abundance in CST III, indicating its role as a transitional species within the vaginal microbiota. Due to its ability to colonize the vaginal niche following microbial disturbances, *L. iners* may provide limited protection against vaginal dysbiosis. However, it has also been implicated in increased susceptibility to BV, sexually transmitted infections (STIs), and adverse pregnancy outcomes, suggesting a dual role as both a symbiont and an opportunistic pathogen ^9,35^. Notably, studies have suggested that *L. iners* can persist in dysbiotic environments and facilitate microbial shifts that may lead to recurrent infections ^36^.

In contrast, 6 of 20 participants exhibited microbial communities similar to CST IV, characterized by reduced *Lactobacillus* dominance, increased bacterial diversity, and enrichment of *Dialister*, *Mageeibacillus* and *Megasphaera*, taxa associated with dysbiotic conditions such as bacterial vaginosis (BV) ^37^. Previous studies have linked CST IV with increased susceptibility to vaginal inflammation and adverse gynecological outcomes, with *Dialister* and *Megasphaera* contributing to BV pathogenesis through the production of biogenic amines, which can alter vaginal homeostasis and promote microbial imbalances ^38^. In addition, the prevalence of CST IV varies among ethnicities and geographic regions, with a higher presence observed in women of African descent populations ^39^. This variation may be influenced by genetic, environmental, and behavioral factors, which warrant further investigation in diverse populations to better understand the determinants of vaginal microbiota composition and stability.

Our study provides the first metataxonomic level characterization of vaginal microbiota in women residing in Chile, offering a foundation for future research on regional microbiome variations and their implications for women’s health. The ability of ONT sequencing to resolve species-level taxonomic assignments with high accuracy offers promising applications for diagnostics and microbiome-based therapeutic strategies. However, limitations such as sequencing error rates and the need for optimized bioinformatics pipelines should be addressed to further enhance its reliability. Future studies with larger cohorts and longitudinal sampling will be essential to elucidate temporal microbiota dynamics, Chilean woman associated microbial profiles and their relationship with health outcomes.

## Data availability

All the data used in this study are available from the cited literature and corresponding websites. All the sequencing data obtained in this study was deposited in the NCBI SRA repository under BioProject code PRJNA1232418.

## Acknowledgements

We thank Dr. Carlos Blondel and the Blondel Lab for their support, including access to laboratory equipment and guidance in molecular biology. We also thank the BioNanotechnology & Microbiology Lab for providing additional laboratory resources. We would also like to thank the 20 volunteers who generously participated in this study by providing samples. This study was partially supported by the Agencia Nacional de Investigación y Desarrollo (ANID) of Chile through various grants: Fondecyt Regular 1221209 and Anillo ATE220061 to J.A.U, and National Doctorate Scholarship 21240354 to P.V.

## Author contributions statement

B.O. conceived and designed the study. B.O. was responsible for participant selection and sample collection. DNA extraction and sequencing were performed by B.O., with contributions from P.V. P.V. conducted the bioinformatic analyses and data visualization. J.A.U. provided conceptual guidance and supervised the entire research process. B.O. and P.V. drafted the initial manuscript, which was subsequently reviewed and refined by J.A.U. All authors reviewed and approved the final manuscript.

## Additional information

### Accession codes

All the sequencing data obtained in this study was deposited in the NCBI SRA repository under BioProject code PRJNA1232418.

## References

1. Tachedjian, G., Aldunate, M., Bradshaw, C. S. & Cone, R. A. The role of lactic acid production by probiotic Lactobacillus species in vaginal health. Res. Microbiol. 168, 782–792 (2017).

2. Pendharkar, S., Skafte-Holm, A., Simsek, G. & Haahr, T. Lactobacilli and their probiotic effects in the vagina of reproductive age women. Microorganisms 11, (2023).

3. Wu, S., Hugerth, L. W., Schuppe-Koistinen, I. & Du, J. The right bug in the right place: opportunities for bacterial vaginosis treatment. NPJ Biofilms Microbiomes 8, 34 (2022).

4. Hou, K. et al. Microbiota in health and diseases. Signal Transduct. Target. Ther. 7, 135 (2022).

5. Workowski, K. A. et al. Sexually Transmitted Infections Treatment Guidelines, 2021. {Centers for Disease Control and Prevention}; https://www.cdc.gov/std/treatment-guidelines/STI-Guidelines-2021.pdf (2021).

6. McKloud, E., et al. Recurrent vulvovaginal candidiasis: A dynamic interkingdom biofilm disease of Candida and Lactobacillus. mSystems 6, e0062221 (2021).

7. van der Veer, C., Bruisten, S. M., van der Helm, J. J., de Vries, H. J. C. & van Houdt, R. The cervicovaginal Microbiota in women notified for chlamydia trachomatis infection: A case-control study at the sexually transmitted infection outpatient clinic in Amsterdam, the Netherlands. Clin. Infect. Dis. 64, 24–31 (2017).

8. Gosmann, C. et al. Lactobacillus-deficient cervicovaginal bacterial communities are associated with increased HIV acquisition in young South African women. Immunity 46, 29–37 (2017).

9. Zheng, N., Guo, R., Wang, J., Zhou, W. & Ling, Z. Contribution of Lactobacillus iners to vaginal health and diseases: A systematic review. Front. Cell. Infect. Microbiol. 11, 792787 (2021).

10. Ravel, J. et al. Vaginal microbiome of reproductive-age women. Proc. Natl. Acad. Sci. U. S. A. 108 **Suppl 1**, 4680–4687 (2011).

11. Nonspecific vaginitis. Diagnostic criteria and microbial and epidemiologic associations. Am. J. Med. 74, A28 (1983).

12. Nugent, R. P., Krohn, M. A. & Hillier, S. L. Reliability of diagnosing bacterial vaginosis is improved by a standardized method of gram stain interpretation. J. Clin. Microbiol. 29, 297–301 (1991).

13. Heikema, A. P. et al. Comparison of Illumina versus nanopore 16S rRNA gene sequencing of the human nasal Microbiota. Genes 11, 1105 (2020).

14. Curry, K. D. et al. Emu: Species-level microbial community profiling for full-length nanopore 16S reads. (2021) doi:10.1101/2021.05.02.442339.

15. Wang, Y., Zhao, Y., Bollas, A., Wang, Y. & Au, K. F. Nanopore sequencing technology, bioinformatics and applications. Nat. Biotechnol. 39, 1348–1365 (2021).

16. Quast, C. et al. The SILVA ribosomal RNA gene database project: improved data processing and web-based tools. Nucleic Acids Res. 41, D590–6 (2013).

17. Parks, D. H. et al. GTDB: an ongoing census of bacterial and archaeal diversity through a phylogenetically consistent, rank normalized and complete genome-based taxonomy. Nucleic Acids Res. 50, D785–D794 (2022).

18. Lillo G, E., Lizama I, S., Medel C, J. & Martínez T, M. A. Diagnóstico de vaginosis bacteriana en un consultorio de planificación familiar de la Región Metropolitana, Chile. Rev. Chilena Infectol. 27, (2010).

19. Villaseca, R. et al. Vaginal infections in a family health clinic in the metropolitan region, chile Infecciones vaginales en un Centro de Salud Familiar de la Región Metropolitana, Chile. Rev. Chilena Infectol. 41–47 (2015).

20. Chen, S., Zhou, Y., Chen, Y. & Gu, J. fastp: an ultra-fast all-in-one FASTQ preprocessor. Bioinformatics 34, i884–i890 (2018).

21. Pruitt, K. D., Tatusova, T., Brown, G. R. & Maglott, D. R. NCBI Reference Sequences (RefSeq): current status, new features and genome annotation policy. Nucleic Acids Res 40, D130–5 (2012).

22. Angly, F. E., Willner, D., Rohwer, F., Hugenholtz, P. & Tyson, G. W. Grinder: a versatile amplicon and shotgun sequence simulator. Nucleic Acids Res. 40, e94 (2012).

23. Mesloub, Y., Beury, D., Vandermeeren, F. & Caboche, S. CuReSim-LoRM: A tool to simulate metabarcoding long reads. Int. J. Mol. Sci. 24, 14005 (2023).

24. Curry, K. D. et al. Emu: species-level microbial community profiling of full-length 16S rRNA Oxford Nanopore sequencing data. Nat. Methods 19, 845–853 (2022).

25. Stoddard, S. F., Smith, B. J., Hein, R., Roller, B. R. K. & Schmidt, T. M. rrnDB: improved tools for interpreting rRNA gene abundance in bacteria and archaea and a new foundation for future development. Nucleic Acids Res 43, D593–8 (2015).

26. O’Leary, N. A. et al. Reference sequence (RefSeq) database at NCBI: current status, taxonomic expansion, and functional annotation. Nucleic Acids Res 44, D733–45 (2016).

27. McMurdie, P. J. & Holmes, S. Phyloseq: a bioconductor package for handling and analysis of high-throughput phylogenetic sequence data. Pac. Symp. Biocomput. 235– 246 (2012).

28. Villanueva, R. A. M. & Chen, Z. J. ggplot2: Elegant Graphics for Data Analysis (2nd ed.). Measurement 17, 160–167 (2019).

29. Ma, Z. S. & Li, L. Quantifying the human vaginal community state types (CSTs) with the species specificity index. PeerJ 5, e3366 (2017).

30. Komiya, S. et al. MinION, a portable long-read sequencer, enables rapid vaginal microbiota analysis in a clinical setting. BMC Med. Genomics 15, 68 (2022).

31. Matsuo, Y. et al. Full-length 16S rRNA gene amplicon analysis of human gut microbiota using MinION^TM^ nanopore sequencing confers species-level resolution. BMC Microbiol. 21, 35 (2021).

32. Lüth, T. et al. Improving analysis of the vaginal microbiota of women undergoing assisted reproduction using nanopore sequencing. J. Assist. Reprod. Genet. 39, 2659– 2667 (2022).

33. Zhou, X. et al. Differences in the composition of vaginal microbial communities found in healthy Caucasian and black women. ISME J. 1, 121–133 (2007).

34. Borgdorff, H. et al. The association between ethnicity and vaginal microbiota composition in Amsterdam, the Netherlands. PLoS One 12, e0181135 (2017).

35. Kadogami, D. et al. Impact of Lactobacillus in the uterine microbiota on in vitro fertilization outcomes. J. Reprod. Immunol. 160, 104138 (2023).

36. Sabbatini, S. et al. Lactobacillus iners cell-free supernatant enhances biofilm formation and hyphal/pseudohyphal growth by candida albicans vaginal isolates. Microorganisms 9, 2577 (2021).

37. Srinivasan, S. et al. Bacterial communities in women with bacterial vaginosis: high resolution phylogenetic analyses reveal relationships of microbiota to clinical criteria. PLoS One 7, e37818 (2012).

38. Nelson, T. M. et al. Vaginal biogenic amines: biomarkers of bacterial vaginosis or precursors to vaginal dysbiosis? Front. Physiol. 6, 253 (2015).

39. Fettweis, J. M. et al. The vaginal microbiome and preterm birth. Nat. Med. 25, 1012– 1021 (2019).

